# Hyper-ovulation in an ovarian-specific transgenic mouse model

**DOI:** 10.1101/2025.02.21.639591

**Authors:** Luis C. Muñiz, James Reilly, Melissa Spigelman, Carlos A. Molina

**Author notes:** Correspondence: Carlos A. Molina, PhD, Department of Biology, Montclair State University, 1 Normal Avenue, Montclair, NJ 07043.

## Abstract

Ovulation is a fundamental prerequisite for achieving successful reproduction. In vertebrates, ovulation is controlled by the cyclical action of hormones, particularly the gonadotropins such as follicle stimulating hormone (FSH) and luteinizing hormone (LH). A critical component of the intracellular activity of these two hormones is relayed by the second messenger cAMP. Although it is well established that a family of transcription factors facilitate cAMP mediated gene expression, it remains unknown how these factors directly affect ovulation. In particular, the Inducible cAMP Early Repressor (ICER) has been implicated in the transcriptional repression of FSH inducible genes during folliculogenesis. Here we show, using an ovarian-specific transgenic mouse model that ICER potentiates ovulation. We observed a twofold rate increase in ovulation for transgenic mice when compared to the wild type in response to exogenous gonadotropin treatment. Furthermore, mature cycling transgenic mice display a significantly enhanced ovulation rate compared to the wild-type. The observed changes in ovulation in the transgenic females are accompanied by altered gonadotropins production and gene expression. Most significantly, the expression of inhibin alpha subunit (INHA) was found to be about 5-fold higher in the transgenic mice. These observations may aid in unraveling some of the molecular mechanisms underlying ovulation and be relevant to the development of novel reproductive technologies.

**Summary Sentence:** Generation of an FSH inducible ovarian specific FLAG-ICER-II© Tg Mice results in hyper-ovulation upon gonadotropin stimulation.

## INTRODUCTION

The ovary is a dynamic organ undergoing constant change, serving as the site of gamete (oocyte) production as well as reproductive hormone synthesis and secretion. The ovarian follicle fosters the oocyte in addition to functioning as the major endocrine and reproductive compartment of the ovary. The process of individual follicle maturation requires both the proliferation and differentiation of the follicular cell compartment [1–4]. This balance entails the coordinated expression of specific genes, dependent on the temporal exchange of many extracellular signals such as hormones and growth factors. Granulosa cells, the cells of the follicles, use an array of signaling pathways to interpret these external cues to ultimately control the switching on and off of genes at the appropriate time during follicular maturation. Surges in FSH and estrogen levels stimulate granulosa cells to proliferate, whereas surges in LH levels inhibit cell growth and induce the differentiation into luteal cells. Although it is well established that cAMP signaling plays a pivotal role in gonadotropin regulation of granulosa cells, it is currently unclear how cAMP mediates the contrasting effect of the gonadotropic hormones on granulosa cell growth and differentiation.

Numerous genes expressed in the ovary are regulated by the cAMP pathway as a consequence of gonadotropin signaling [5]. During the rodent estrous cycle, FSH secreted from the anterior pituitary results in the expression of many FSH-responsive genes important for the growth and maturation of the ovarian follicle in the ovary. However in response to the preovulatory LH surge, many FSH-responsive genes are rapidly down-regulated.

The nuclear response to the cAMP pathway is mediated by a large family of transcription factors [6]. The best characterized of these factors are the cAMP-Response Element (CRE)-Binding Protein (CREB), CRE Modulator (CREM) and Activating Transcription Factor 1 (ATF-1). An isoform of CREM that is of particular interest is ICER which has been shown to mediate the antiproliferative activity of cAMP in granulosa cells regulating the expression of INHA, cytochrome P450 (CYP19a) and cyclin D2 [7–9].

CREB and CREM genes encode several nuclear factors that can act as transcriptional activators or repressors of cAMP-responsive genes [6]. These transcription factors exert their effects upon binding to CREs within the promoters of cAMP-responsive genes. ICER is unique among the other CRE binding proteins in that its expression is induced by cAMP from an internal promoter within *Crem* gene [10]; the promoter itself is responsive to cAMP thereby allowing ICER to autoregulate its own expression via negative feedback. ICER shares the DNA binding and dimerization domains with the other CREM isoforms, but lacks the kinase and transactivation domains. ICER therefore functions as a dominant negative transcriptional repressor by binding as a homodimer or heterodimer with other CRE-binding family members.

This unique feature endows ICER with a key role in mediating the repression of cAMP-dependent transcription. In the ovaries of adult cycling rats, *Crem* mRNA levels of ICER isoforms have been shown to be selectively induced in the granulosa cells of preovulatory follicles in response to the ovulatory surge of LH [11]. Similarly, ICER expression was found to be induced in granulosa cells of PMSG-primed immature rats injected with human Chorionic Gonadotropin (hCG), whereas PMSG alone did not induce ICER expression [11]. This induction of ICER in response to hCG has been proposed to mediate the suppression of FSH inducible genes, such as INHA [7,11], CYP19a [8] and cyclin D2 [9].

The anterior pituitary glycoprotein, FSH, is essential for folliculogenesis. This is evident in hypophysectomized rats which lack sustained follicular growth due to the lack of gonadotropins. However, treatment with FSH promotes the formation of large antral follicles [12]. Another major effect of FSH on granulosa cells is to induce the expression of LH receptors and acquire LH responsiveness [13]. FSH has been shown to stimulate cyclin D2 mRNA via a cAMP/PKA pathway in granulosa cells [4,14]. However, a luteinizing dose of LH in hormonally primed hypophysectomized female rats results in a rapid decrease in cyclin D2 mRNA and protein levels [4]. Presently, it is mostly unclear how cAMP mediates the actions of both FSH and LH in producing their contrasting effects on granulosa cell growth and cyclin D2 expression. One possibility might be related to differential expression of the transcription factors that mediates the cAMP pathway in ovarian cells.

Therefore, in order to determine the role of ICER in ovarian function *in vivo,* a mouse model with restrictive expression of ICER to the ovaries, particularly within the granulosa cells, was generated using the 2.5kb fragment of the FSH/PMSG inducible INHA gene promoter previously characterized to drive the expression of the transgene to the granulosa cell compartment of the ovarian follicle [15]. Here we report that overexpression of ICER in the ovaries leads to an increase in the ovulation rate either in the hormone-primed immature or mature cycling females; thereby conferring an important physiological role of ICER in regulating normal ovulation rate.

## MATERIALS AND METHODS

### Animals

FVB mice (Taconic Farms, Germantown, NY) were used in the present studies. Animals were housed and handled in accordance with protocols approved by the Institutional Animal Care and Use Committee (IACUC) of the Rutgers-New Jersey Medical School.

### Generation of transgene cassettes

The mouse ICER-II© cDNA was subcloned into the pCMV2-FLAG to generate an amino terminal tagged ICER II© expression plasmid. The 5’ end of the FLAG-ICER DNA was extended with the 45 bp region of the 5’ UTR of ICER fused to the ATG of FLAG by two rounds of PCR using the following primers: Forward primer 1^st^ Round (5’-CCT GCA GTG GAC TGT GGT ACG GCC AAT AAG ACC ACT CTA TAT GC-3’), Forward primer 2^nd^ Round with *PstI* site (5’-ACC ACT CTA TAT GCA AAA GCC CAA CAT GGA CTC CAA AGA C-3’) with Reverse primer containing an *XbaI* site (5’-CGT CTA GAT ACT AAT CTG TTT TGG GAG-3’). The final PCR product was subcloned into PCR 2.1 TOPO vector. The SV40 intron and ploy A site was PCR amplified from pGL2-Basic using the following primers: Forward (5’-CCG TCT AGA AAT GTA ACT GTA TTC AGC G-3’) and Reverse (5’-CGC TGC AGC TCG AGA GAC ATG ATA AGA TAC ATT G-3’) and subcloned into PCR 2.1 TOPO vector. The UTR-FLAG-ICER II© DNA was excised from TOPO vector as an *XbaI/XbaI* fragment and inserted in the *XbaI* site upstream of the SV40 intron poly (A) signal of the SV40 TOPO plasmid. The resulting construct was then digested with *PstI* to cut out the 1.4kb fragment. The 3’ over-hang ends were blunt ended with T4 DNA polymerase and subsequently subcloned into the *EcoRV* site of Bluescript (SK-) downstream of the 2.5 kb mouse inhibin alpha promoter [15]. For the production of Tg mice, the transgene cassettes were isolated by excising the *XhoI* fragment from the resulting Bluescript (SK-) final construct and was microinjected into fertilized oocytes of FVB x FVB mice. The injected oocytes were then implanted into pseudopregnant CD-1 females (Charles River).

### Genotyping

Genomic DNA was extracted from tail biopsies obtained from 2 to 3 week old mice using the phenol:chloroform method as described by The Jackson Laboratory. Genomic DNA was digested for 4 hr with *BamHI* and *XbaI*. After digestion, DNA was fractionated by electrophoresis in 0.8% agarose gel, denatured and transferred to a nylon membrane (Hybond-N+, GE Healthcare). Radiolabeled DNA fragment comprising the FLAG-ICER region of the transgene was used as hybridization probe. Hybridization was done overnight at 65°C in hybridization solution (5X SSC, 5X Denhardt’s reagent, 0.5% SDS) including 200 ⎧g/ml denatured salmon sperm DNA. The membrane was washed twice with 2X SSC, 0.1% SDS at 65°C for 30 min and twice with 1X SSC, 0.1% SDS at 65°C for 30 min. Membranes were exposed to films for autoradiography.

### RNase Protection Assay

RNA was extracted from primary granulosa cells or ovarian tissues using TRIZOL Reagent (Invitrogen) according to the manufacture’s instructions. Aliquots of 5 µg of total RNA were subjected to RNase protection analysis essentially as described [10]. Briefly, RNase Protection assays were performed using the RPA III Ribonuclease Protection Assay kit (Ambion) following the manufacturer’s instructions using radioactive riboprobes that recognizes either the exogenous or endogenous ICER transcripts. A mouse ®-actin probe was used for loading control where indicated. Samples were separated on a denaturing polyacrylamide gel (6% polyacrylamide/ 8 M urea) and the gels were vacuum dried at 80°C and visualized by autoradiography.

### [〈-^32^P] UTP Labeling of RNA Probe

The p6N/1 template DNA used to detect exogenous ICER and the p75 template DNA used to detect endogenous ICER have been previously described [10]. For loading control, pTRI-®-actin-mouse antisense control templates (Ambion, Austin, TX) were used. The RNA probes used in the RNase protection assay were generated using MAXIscript In vitro Transcription Kit (Ambion) following the manufacturer’s instructions. The probes were purified and free nucleotides were removed by column purification using Probe Quant G-50 Micro-column.

### Antibodies

The anti-ICER polyclonal antibody was raised against bacterially purified ICER-IIγ and previously characterized [10]. This antibody has been shown to cross react with other CREM isoforms and ubiquitinated forms of ICER and does not cross react with CREB. Anti α-tubulin antibody (polyclonal), anti-FLAG M2-Peroxidase (HRP) antibody (monoclonal) were used for Western blot analysis and Anti-FLAG M2 antibody (monoclonal) was used for immunohistochemistry and purchased from Sigma, St. Louis, MO. All secondary antibodies, goat anti-rabbit and goat anti-mouse horseradish peroxidase conjugated, were purchased from Jackson ImmunoReseach Laboratories, Inc. West Grove, PA. *Immunoblotting*.

Whole cell protein lysates obtained using Trizol reagent were boiled and protein quantification was performed using Bio-RAD DC reagent with BSA as a standard. Equal amounts of protein (20-30 ⎧g) were mixed with 2X loading buffer (20% glycerol, 0.1% bromophenol blue, 2mM DTT). Samples were boiled for 5 min and loaded onto 15% SDS-polyacrylamide gels. The gels were run at 200 volts for 55 min. The proteins were transferred onto PVDF membrane for 80 min at 100 volts. The membranes were blocked for 10 min in 1X PBS containing 5% non-fat milk or BSA (Following the recommendations of the manufacturer of the antibody being used) and probed overnight at 4°C with either anti-FLAG M2-peroxidase (HRP) monoclonal antibody (Sigma 1:500) or with a previously characterized rabbit polyclonal anti-ICER antibody (1:1000). Blots were washed in 1X PBS with 0.1% Tween-20 four times for 15 min each, then probed with anti-rabbit horseradish peroxidase conjugated antibody at a 1:20,000 dilution in 5% non-fat milk for 45 min, washed 4 times for 4 min each in 1X PBS with 0.1% Tween-20 and visualized using enhanced chemiluminescence and autoradiography.

### Immunohistochemistry

Ovaries were fixed in 10% formalin for 24 hr, dehydrated, embedded in paraffin and sectioned at 5 µm. Standard hematoxylin and eosin staining was also preformed. Briefly, sections were baked in an oven at 56°C for 30 min, followed by deparafinization in Xylene. Sections were then rehydrated in a series of sequential ethanol baths ranging from 100%, 95%, 75% and 50%. Sections were rinsed and subjected to peroxidase quenching. An antigen retrieval step was preformed using 0.01M Citrate Buffer solution (pH 8.0). Sections were either incubated with a 1:5,000 dilution of a rabbit polyclonal antibody that recognizes ICER or a 1:2,500 dilution of a mouse monoclonal antibody that recognizes FLAG overnight at 4°C. Sections were then washed and incubated with a biotinylated goat anti-rabbit antibody or biotinylated goat anti-mouse antibody (Zymed Laboratories Inc.) for 10 min at room temperature. Sections were then washed, incubated for 10 min at room temperature with a horseradish peroxidase-conjugated streptavidin (Zymed Laboratories Inc.), washed, and incubated with a Subrate-Chromogen Mixture (Zymed Laboratories Inc.) for 10 min at room temperature for antigen detection (brown reaction product). Sections were counterstained with hematoxylin and analyzed by light microscopy.

### Gonadotropin Treatment and Oocyte harvesting

Intraperitoneal (IP) gonadotropin administrations were performed as described [14]. Mice were subsequently sacrificed at various time points after PMSG or hCG treatment and the ovaries were removed and either immediately frozen in liquid nitrogen and stored at −80°C until further processed or immediately fixed in 10% formalin.

Immature FVB females (3-4 weeks) were injected with 5 IU of PMSG (at noon) followed 48 h later with 5 IU hCG and mated with FVB stud males. The next morning, upon detection of a vaginal plug, the fertilized oocytes were collected from the oviduct in M2 medium. The embryos were then incubated in M2 with hyalurinadase (300 ⎧g/ml) to detach the follicle cells from the embryos, washed several time with M2 media and incubated in M16 medium at 37°C in 5% CO_2_. The fertilized oocytes were cultured for 4 days and allowed to develop to the blastocyst stage. After 4 days the total number of blastocysts were determined and a percentage from the total number of embryo were calculated.

Determination of oocyte release in mature, 2-month old, wild-type or transgenic mice was preformed by introducing a stud male. Female mice were checked for the presence of a vaginal plug every morning as indirect indication of ovulation, subsequently sacrificed and fertilized embryos collected as described above.

### In Vitro Ooctye Maturation

Immature FVB females (3-4 weeks) were injected IP with 5IU of PMSG (at noon) followed 48h later with 5IU hCG and mated with FVB stud males. The next morning the females were checked for the presence of vaginal plugs as an indicator of successful mating. The ovaries and the fertilized oocytes were collected from the oviduct in M2 medium. The embryos were then incubated in M2 with hyalurinadase (300⎧g/ml) to detach the follicle cells from the embryos, washed several time with M2 media and incubated in M16 medium at 37°C in 5% CO_2_. To assess the viability of the collected embryos, the fertilized oocytes were cultured for 4 days and allowed to develop to the blastocyst stage. After 4 days the total number of blastocysts were determined and a percentage from the total number of embryo were calculated.

### Vaginal Cytology

Changes in vaginal epithelial cells were monitored by taking daily vaginal smears using 3-month-old transgenic or wild type mice as described by Cooper et al., [16]. Vaginal smears were obtained daily at 10AM and 5PM for two week so that the cycling pattern could be determined accurately. Loose epithelial cells were obtained by gently inserting a sterile micropipette tip no more then 1mm and expelling 20μl of PBS into the vaginal cavity and aspirating it back into the tip twice. The collected fluid was dispensed evenly onto a microscope slide and the smear was viewed immediately under low magnification (10X) for the presence of leukocytes, nucleated epithelial cells or cornified cells. The vaginal smears were classified as diestrus (presence of leukocytes), proestrus (presence of nucleated epithelial cells), estrus (presence of cornified epithelial cells) or metestrus (presence of cornified epithelial cells and leukocytes) as characterized in earlier report [16]. Slides were read and recorded independently by three different people and a continuous record was kept on each female. Mice were sacrificed and ovaries removed towards the end of the two-week study, obtaining a representative coverage of the estrous cycle. Ovaries were frozen rapidly on dry ice and stored at −80°C until use. Blood was collected for gonadotropin measurements by cardiac puncturing.

### Hormone Assays

LH and FSH were measured by mouse immunoradiometric assay (IRMA) and mouse RIA, respectively, by the Ligand Assay and Analysis Core Laboratory, Center for Research in Reproduction, University of Virginia, Charlottesville. The sensitivity of the IRMA for LH was 0.07 ng/ml (average intra-assay and inter-assay coefficients of variation for the Quality Controls are 3.6% and 11.1%) and the sensitivity of the RIA for FSH was 2 ng/ml with less than 0.5% cross-reactivity with other pituitary hormones (intra-assay variation, 10%; inter-assay variation, 13.3%).

Progesterone levels were measured in serum, according to the manufacturer’s instructions, using a direct solid-phase enzyme-immunoassay (DRG Progesterone ELISA kit). Before the assay, the serum samples were extracted with 6.6 vol of ethyl ether; the extracts were dried and reconstituted in zero standard control serum provided by the manufacturer. Sera from WT and ICER transgenic mice were assayed in duplicate in the same assay. Progesterone concentration measured by a spectrophotometer at 450 nm, against a standard curve.

### RNA Extraction, Microarray Hybridization and Data Analysis

Total RNA from the ovaries of immature mice subjected to exogenous gonadotropin regimen were isolated by using the method of Trizol (Invitrogen, Carlsbad, CA) extraction coupled in-column DNase digestion using the RNeasy Midi kit from Qiagen (Valencia, CA) according to the manufacturer’s protocol.

After RNA isolation, all the subsequent technical procedures, including quality control, concentration measurement of RNA, cDNA synthesis, and biotin labeling of cRNA, hybridization, and scanning of the arrays, were done at CINJ Core Expression Array Facility of Robert Wood Johnson Medical School (New Brunswick, NJ). Affymetrix mouse genome 430A 2.0 array containing over 22,600 probe sets representing transcripts and variants from over 14,000 well-characterized mouse genes was used to probe the gene expression profile in transgenic mice. Each array was hybridized with cRNA derived from the total RNA sample from either transgenic or wild-type mice (total 2 chips were used in this study). After hybridization and washing, the intensities of the fluorescence of the array chips were measured by the Affymetrix GeneChip Scanner. The expression analysis file created from each sample (chip) scanning was imported into GeneSpring 6.1 software (Silicon Genetics, Redwood City, CA) for further data characterization. Lists of genes that were either induced or suppressed >2-fold between transgenic and wild-type mice are reported.

### Quantitative Real-time PCR

As a validation of the microarray results, real-time RT-PCR analysis of several genes was performed. Total RNA isolated from ovaries used for DNA microarray analysis was utilized. One microgram of total RNA was reversed transcribed using advantage RT-for-PCR kit (BD Clontech) following the manufacture’s protocol. Briefly, the RNA and oligo(dT)18 mix was denatured at 72°C for 10 min then 200 U of Moloney murine leukemia virus reverse transcriptase enzyme were used in a final reaction of 20⎧l. cDNA was diluted with 30⎧l of water. One microliter of the diluted cDNA was used for real-time PCR using the fluorescent dye SYBR Green to monitor DNA synthesis (SYBR Green PCR Master Mix, Applied Biosystems) using specific primers designed for mice, Cyp17a1, Cyp19a1, cyclin D2, Des, INHA and Gapdh. The following primer sets were used:

Gapdh 5’-ATGTTCCAGTATGACTCCACTCACG-3’, 5’-GAAGACACCAGTAGACTCCACGACA-3’;

Cyp19a1 5’-CCTGACACCATGTCGGTCACT-3’, 5’- GGGCTTAGGGAAGTACTCGAG-3’;

Cyp17a1 5’-GAAGTGCTCGTGAAGAAGGG-3’, 5’- CCCAGGACATCCACAATACC-3’;

Cyp17a1 5’-GAAGTGCTCGTGAAGAAGGG-3’, 5’- CCCAGGACATCCACAATACC-3’;

Des 5’-ACTCCCTGATGAGGCAGAT-3’, 5’-GCTGACAACCTCTCCATCCCG-3’;

Cyclin D2 5’-GTTCTGCAGAACCTGTTGAC-3’, 5’- ACAGCTTCTCCTTTTGCTGG-3’;

INHA 5’- CTGTGCCTGTGTCCTTGGTA-3’, 5’- CCAGGAAAGGAGTGGTCTCA-3’

Each PCR reaction was performed in triplicate. Samples were compared by the relative (comparative) threshold cycle (Ct) method. Fold induction or repression was measured relative to controls and calculated after adjusting for GADPH by using 2-ΔΔCt, where ΔCt = Ct for gene of interest - Ct GADPH and ΔΔCt = Ct transgenic - Ct wild-type. Data are presented as the relative average value of the gene investigated and normalized with the average value of the housekeeping gene (Gapdh). Melting curves were analyzed to confirm amplification specificity. *Statistical Analysis.* Differences between treatments were analyzed for significance by Student’s *t*-test.

## RESULTS

### Generation of an ovarian specific transgenic of ICER

Numerous genes expressed in the ovary are regulated by the cAMP pathway as a consequence of gonadotropin signaling [5]. During the rodent estrous cycle, FSH secreted from the anterior pituitary results in the expression of many FSH-responsive genes important for the growth and maturation of the ovarian follicle in the ovary. However in response to the preovulatory LH surge, many FSH-responsive genes are rapidly down-regulated. ICER has been implicated in the transcriptional repression of FSH inducible genes during folliculogenesis [7–9,11]. Yet no *in vivo* model systems to date are available to further elucidate ICER role in ovarian function. We therefore sought to generate ovarian specific ICER transgenic mice.

The DNA construct used to generate the transgenic mice contained FLAG-ICER cDNA sequence, flanked by 48 base pairs derived from the 5’ UTR of the endogenous ICER RNA, under the control of a 2.5 kb fragment of the FSH/PMSG inducible inhibin alpha-subunit gene promoter (Figure 1A) to restrict the expression of the transgene to the granulosa cell compartment of the ovarian follicle as previously demonstrated [15]. After the DNA construct microinjection into fertilized oocytes from FVB mouse strain three founders were identified by Southern blot. Genotyping was performed using genomic DNA extracted from mouse-tail biopsies of 2-week-old pups. The extracted DNA was analyzed by Southern blot using a probe against FLAG-ICER to detect the 2.18 kb BamHI-XhoI fragment generated from the integrated cassette. The probe would also detect three endogenous CREM gene fragments, at 522bp, 8.5kb and 18.5kb from the resulting digestion (Figure 1B). We identified three independent transgenic founder mice TGN1, TGN30 and TGN59 (Figure 1C). At eight weeks of age, the three transgenic founder mice were backcrossed to FVB female mice. Two of the lines (TGN1 and TGN59) transmitted the transgene to the F1 generation and were phenotypically indistinguishable from each other and from their wild-type littermates.

**Figure 1.**
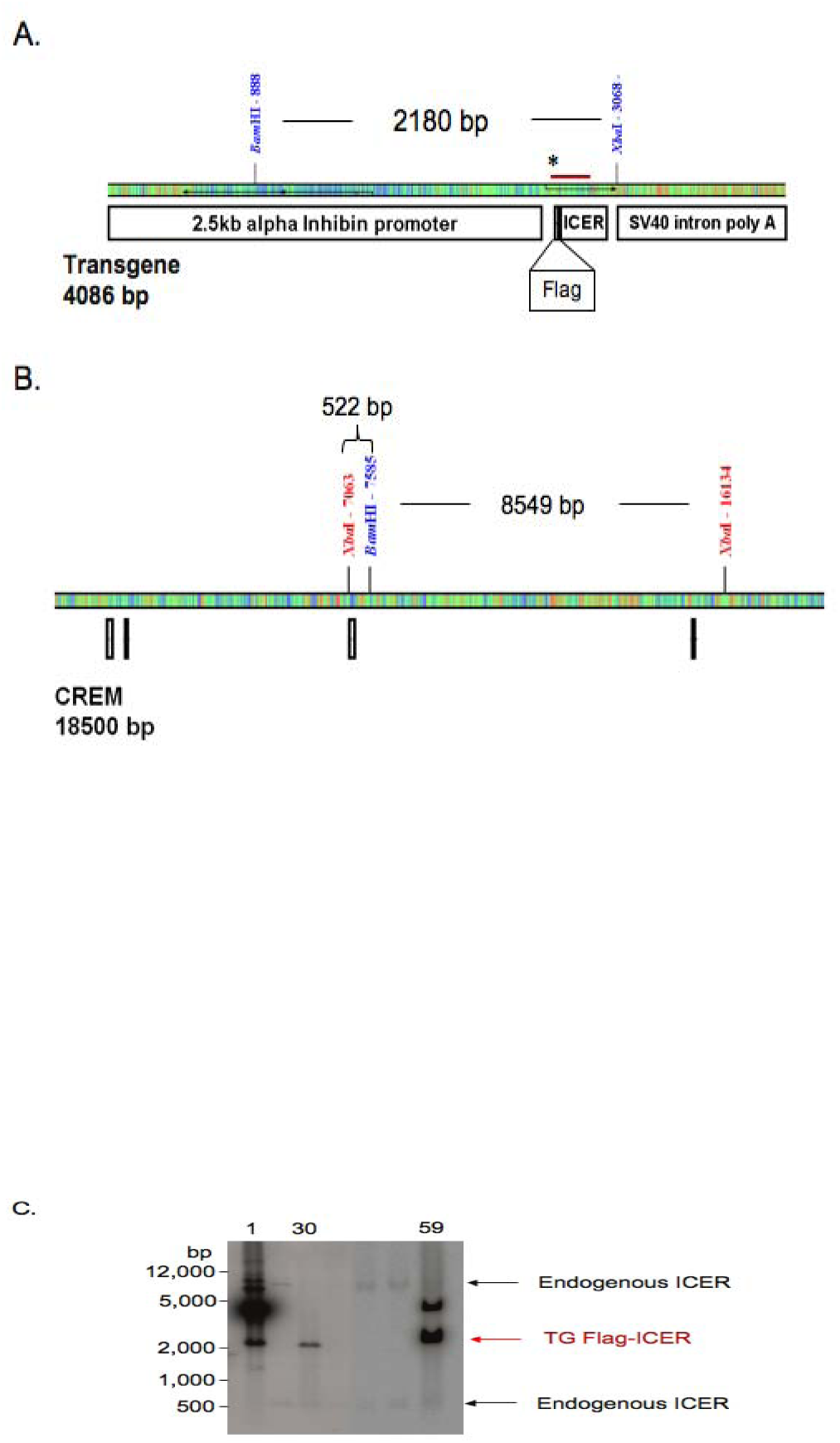
ICER ovarian-specific transgenic mice. Genomic DNA was digested with *BamH* I and *Xba* I and analyzed by Southern Blot using a probed against FLAG-ICER. **A**. Linear representation of the transgene digested with *BamH I* and *Xba I* resulting in a 2.18kb fragment. The red line represents the area used to generate the FLAG-ICER probe. **B.** Diagram representing the results of a sequence alignment between the FLAG-ICER cDNA against the CREM gene. The boxed region represent the location of the ICER-II_©_ exons with the 18.5kb portion of the of the CREM gene. Labeled are the predicted fragments that hybridize with FLAG-ICER probe. **C.** Autoradiograph representing Southern blot analysis on genomic DNA extracted from F0 pups. Labeled are the ID numbers of positive transgenic founder mice.

### Characterization of transgenic mice lines

Ovarian specific expression of FLAG-ICER was confirmed by Western blot analysis using lysates prepared from different tissues of mature TGN1 mice. Figure 2A shows that FLAG-tagged ICER was specifically expressed in the ovary when protein lysates were probed with antibodies raised against ICER or those specific for FLAG peptide. Similar results were obtained with TGN59 (not shown). In order to assess the ovarian responsiveness of the transgenic mice to exogenous gonadotropin treatment, immature 3-week old mice were injected with either PMSG alone for 48hr or followed by hCG for 24hr post PMSG. RNA and protein levels of exogenous ICER were induced in the ovaries of transgenic mice TGN1 48hr after PSMG treatment (Figure 2 C-D). Enhanced expression of the transgene is detected in the transgenic mice 24 hr after hCG treatment when compared to PMSG treatment alone. Similar results were also obtained with TGN59 (not shown).

**Figure 2.**
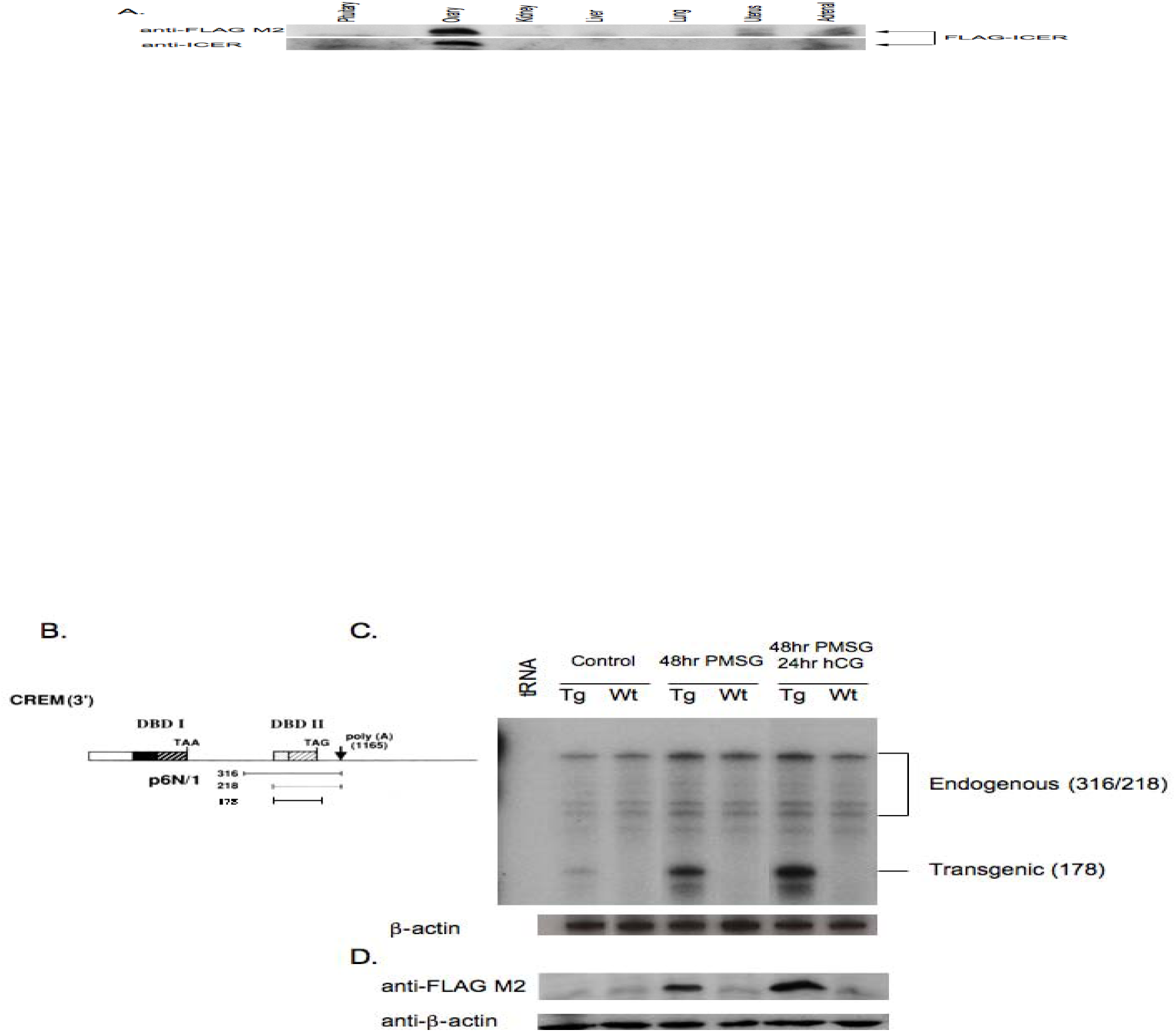
Characterization of an ovarian specific ICER transgenic mice. **A.** Immunoblot analysis of transgene expression in different tissue lysates from mature mice probed with either anti-Flag M2 or anti-ICER antibodies. **B.** Schematic representation of the 3’ end of the CREM depicting the exons coding for DNA Binding Domains I and II (DBD I and DBD II). Under the CREM gene are the predicted protected fragments produced using the probe (p6N/1) in RNase protection assays. Since FLAG-ICER consists of the cDNA of ICER-II© it would only produce a protected fragment of 178 bp. Endogenous CREM isoforms will produce protected fragments of 316 and 218 bps. **C.** Ribonuclease protection analysis of FLAG-ICER mRNA during exogenous gonadotropin treatment in immature transgenic and wild-type female mice. For loading control, ®-actin mRNA levels were determined. **D.** Immunoblot analysis of protein fraction from samples in B probed with either anti-Flag M2 or anti-ICER antibodies.

The localization of the induced transgene expression within the ovarian compartment was determined by immunohistochemical analysis. PMSG-primed immature female mice were injected with hCG and their ovaries were collected 24hr later and sectioned for analysis. We observed that exogenous gonadotropin treatment resulted in tissue specific expression of FLAG-ICER within the granulosa cells of the preovulatory follicle and in the luteinizing granulosa cells of the developing corpus luteum in tissues probed either with anti-ICER or anti-FLAG M2 antibody (Figure 3 A-B). These results were consistent in both transgenic lines (not shown). This data collectively demonstrates that we have successfully generated an animal model where the expression of ICER is induced with PMSG prior to hCG stimulation in granulosa cells. Furthermore, we have achieved an enhanced expression beyond 12 hr of hCG treatment. This mouse model clearly manifests alterations in the temporal expression pattern of ICER in the ovaries, contrasting with the normally occurred expression of endogenous ICER in response to exogenous gonadotropins [11].

**Figure 3.**
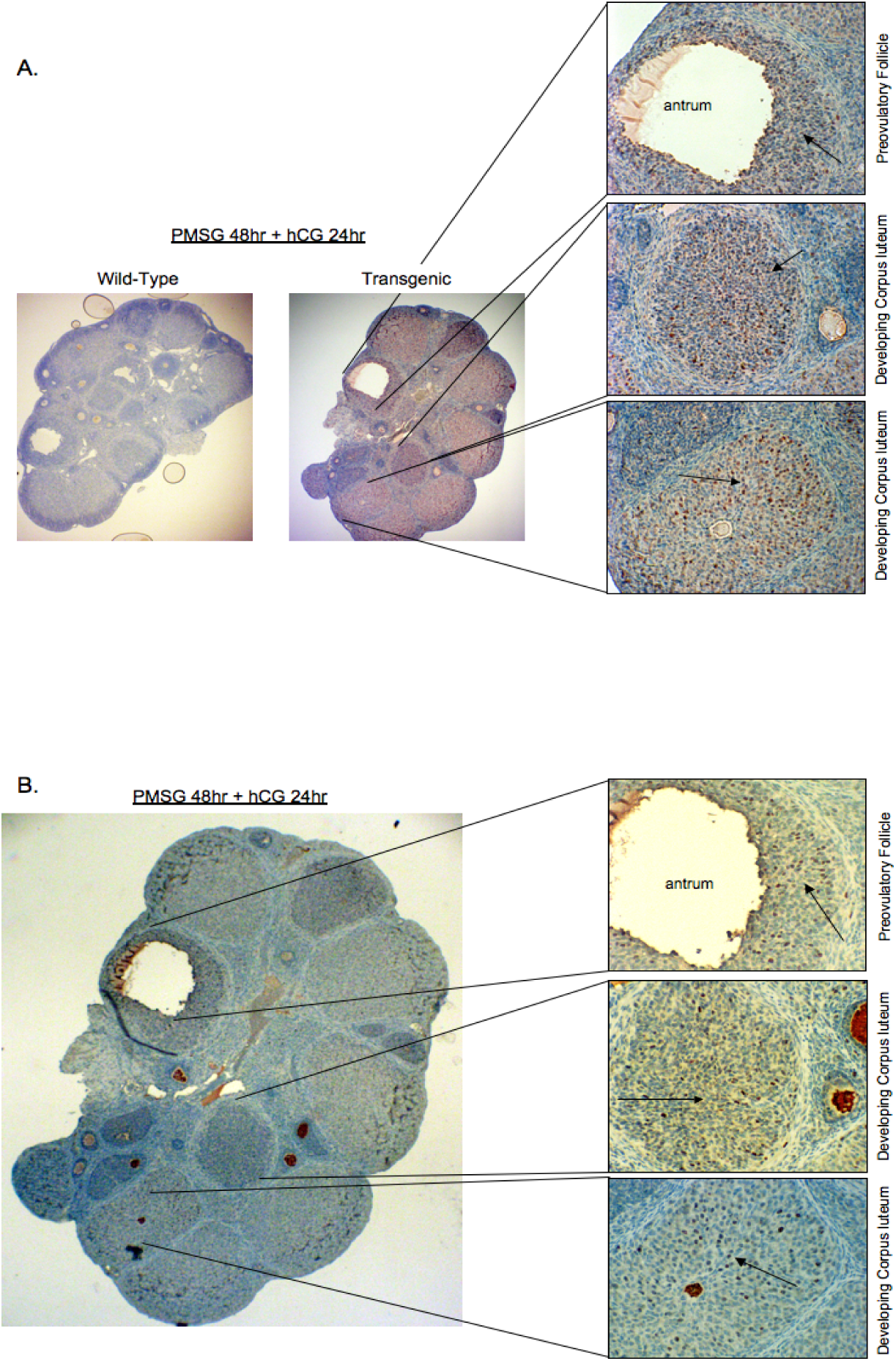
Granulosa cell specific localization of ICER exogenous expression in transgenic mice. **A-B.** Ovary tissue sections from transgenic mice or wild-type littermate primed with PMSG for 48 hrs followed with hCG treatment for an additional 24 hrs and analyzed by immunohistochemistry using (**A**) anti-ICER antibody or (**B**) anti-FLAG M2 antibody. Arrows indicated region with brown staining indicating FLAG-ICER expression. Right hand panel represent 40X magnification of tissue sections. On the left hand panel, 100X magnification of tissue section and the location of the area of detail are indicated.

### Phenotypic characterization of the ovarian specific ICER transgenic mice

In order to determine the physiological effect of FLAG-ICER over-expression on ovarian function, in particular on ovulation, we compared the numbers of released oocytes between immature transgenic and wild type mice. Flag-tagged ICER transgenic mice displayed hypersensitivity to the ovulatory effects of PMSG and hCG, resulting in a two-fold increase in released oocytes compared to wild type littermates (Figure 4A and Table 1). However, we did not observe an increase in litter size between transgenic and wild-type female mice. This apparent discrepancy is most likely related to the probability that a limited number of blastocysts implantation could be a determining factor. In order to rule out intrinsic abnormalities due to ICER overexpresion or as a result of hyperovulation, oocyte viability was assessed. The harvested oocytes were subjected to *in vitro* culture and maturation. The percentage of oocytes proceeding to blastocysts as seen after 4 days in culture were found to be comparable between transgenic and wild type mice (Table 1), suggesting that implantation rather then maturation would account the lack of increase litter size in the transgenic female mice.

**Figure 4.**
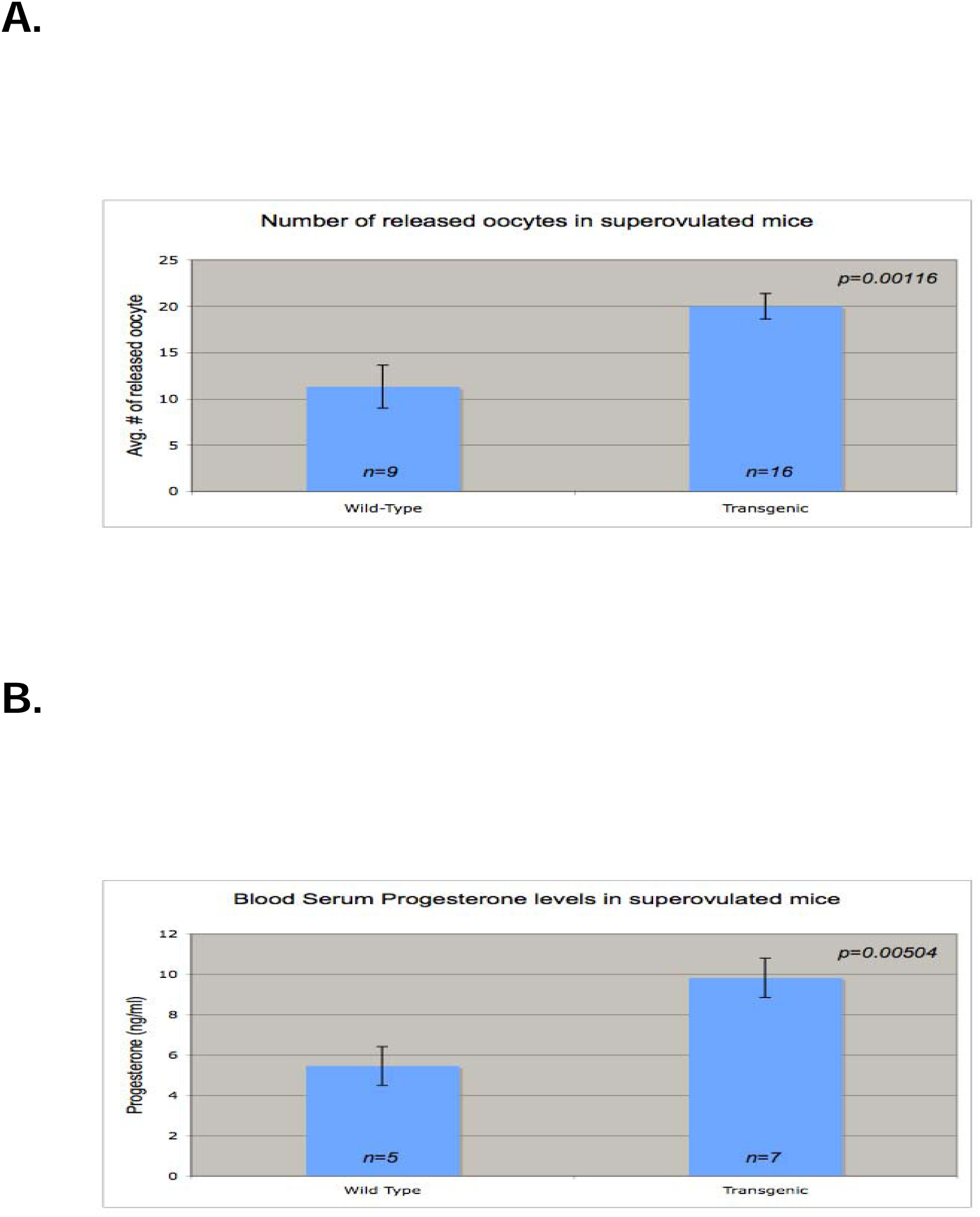
Hyperovulation and high progesterone levels in immature transgenic mice. A. Number of released oocytes from gonadotropin-treated transgenic or wild type mice. B. Mean concentration of progesterone in blood serum from gonadotropin-treated wild-type or transgenic mice. Error bars are s.e.m. *P* values calculated using a one-tailed Student’s *t*-test.

**TABLE 1.** *In vitro* culture of oocytes.

|  | Control | Transgenic |
| --- | --- | --- |
| Number of oocytes | 53* | 144* |
|  | (n=5)† | (n=7)† |
| Blastocysts¶ (%) | 83 | 82.6 |

Sexually immature, control (wild-type) and Flag ICER transgenic female were treated with gonadotropins and mated with wild-type males.
\* Total number of oocytes isolated and used for *in vitro* culture
† Number of females from which oocytes were isolated.
¶ Percentage of fertilized oocytes that formed blastocysts.

In light of our observation, we next measured serum progesterone levels to determine whether the increased number of released oocytes correlates with an increased number of developing corpus luteum. We found that transgenic mice exhibited twice the serum progesterone level as wild type mice (Figure 4B). This observation complements and supports the two-fold rate increase in ovulation observed in the transgenic mice.

### Ovulation rate during the estrous cycle in transgenic mice

Since we had seen an increase rate of released oocytes in gonadotropin stimulated immature mice, we next wanted to determine if mature transgenic mice release more oocytes then wild-type mice. In this experiment, two-month old female mice were mated with wild-type male FVB mice and sacrificed upon the detection of a vaginal plug, during which time the ovaries and oocytes were collected. Figure 5A shows that mature transgenic females ovulate (11.75 ± 0.46; n=12) significantly more then wild-type littermates (8.67 ± 0.21; n=6). To determine if the observed rate of ovulation correlates with changes in ovarian weight due to the consequent rise in corpora lutea formation, ovarian weights were assessed. We found that transgenic ovaries weighed (5.51 ± 0.31 mg; n=6) significantly more then wild-type (3.38 ± 0.53 mg; n=3) (Figure 5B). Collectively, increased rate of ovulation in these animals correlates with an increase in the number of CL in the ovaries, which is reflected by an increase in ovarian weight.

**Figure 5.**
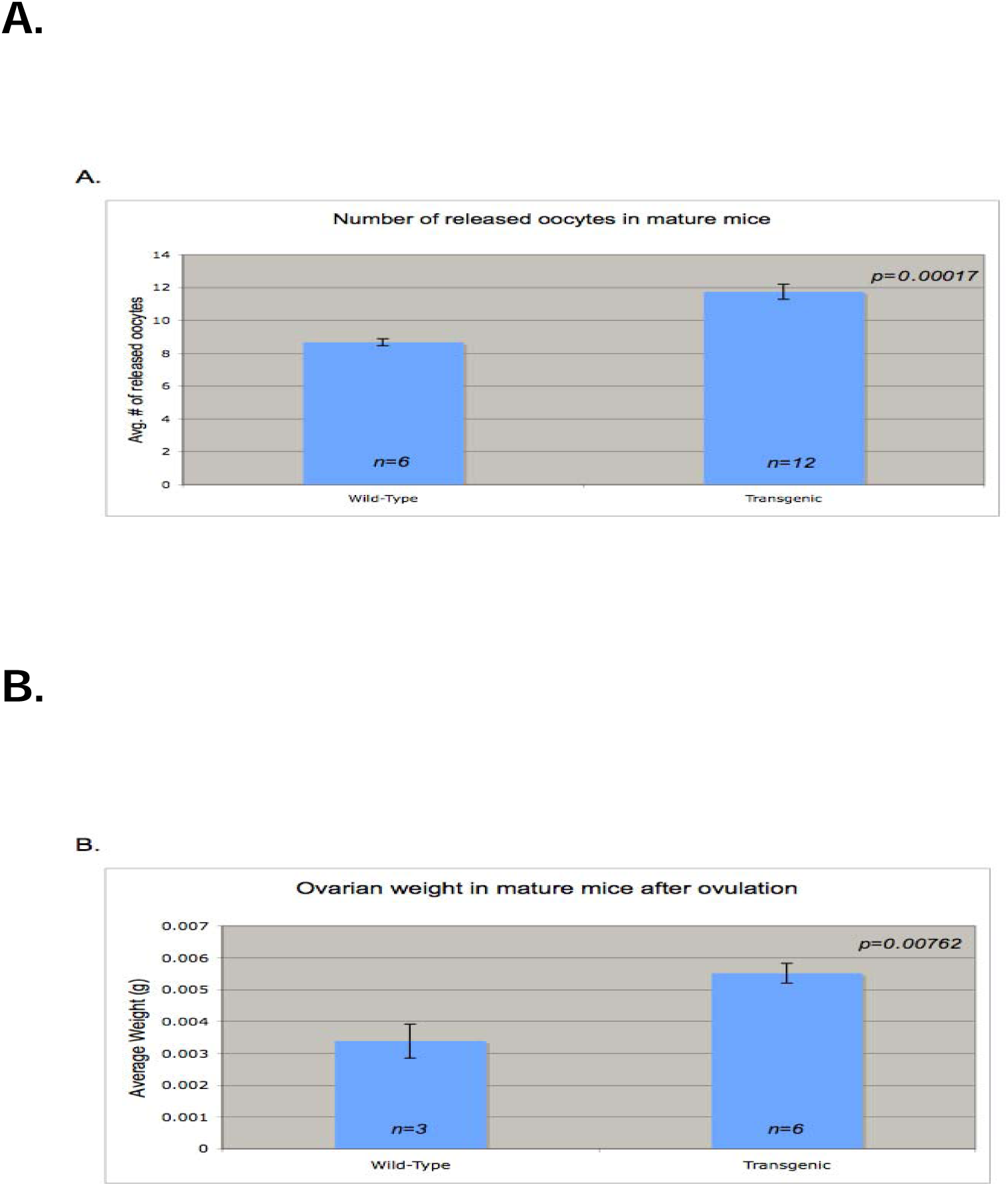
High ovulation rate in Mature Transgenic Mice. A. Mean number of released oocytes from mature two month old transgenic or wild type mice. B. Mean weight of ovaries after ovulation. Error bars indicate s.e.m. and *P* values calculated using a one-tailed Student’s *t*-test in A-B.

### Transgene expression during the estrous cycle

Exogenous gonadotropin-treatment of immature mice serves as a well-known model for events that occur during the normal reproductive cycle in mature females. However, we next wanted to assess the induction of Flag-tagged-ICER in mature estrous cycling mice. For this purpose, RNA and protein were isolated from the ovaries of three-month old estrus cycling transgenic mice at different stages of the cycle that was determined by daily examination of vaginal cytology. Blood serum was simultaneously obtained for hormonal analysis. Figure 6A is representative data of a Ribonuclease Protection Assay using a riboprobe against ICER. The data depicts a biphasic RNA profile of the transgene expression. The transgene is apparently induced during metestrus with a stronger induction during early diestrus and with levels dropping late into diestrus. A rise in the transgene is seen again during late proestrus with a substantial rise in early estrus and subsequent drop throughout estrus. Western blot analysis on the protein samples from these ovaries using an anti-FLAG M2 antibody correlate with the RNA levels, such that the expression levels of the transgene in induced during diestrus and within late proestrus / early estrus (Figure 6B). Endogenous ICER mRNA levels have been reported to be induced during late proestrus in response to LH [11]. We show similar induction in the ICER transgenic mice, however we also detect inducibility early during the cycle during diestrus. This data clearly demonstrates an altered temporal expression pattern of ICER in our transgenic mice.

**Figure 6.**
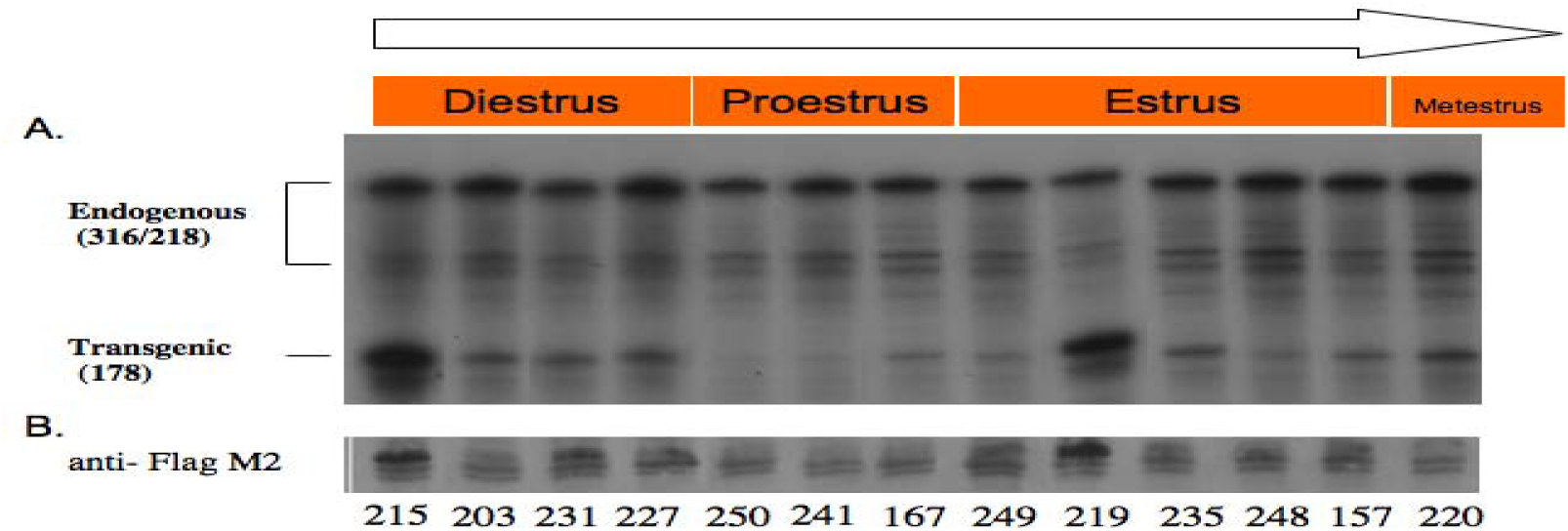
Transgene expression during the estrous cycle. Stages of the Estrous cycle were determined by daily examination of vaginal cytology in three-month old TGN 1 mice. Several animals were sacrificed to cover the different stages of the Estrous cycle. **A.** Ribonuclease protection analysis of FLAG-ICER mRNA using the probe (p6N/1) to analyze the transcriptional levels of FLAG-ICER mRNA. **B.** Immunoblot analysis of protein fraction from samples in (A) probed with anti-FLAG M2 antibody. Numbers correspond to the identification number of each transgenic mouse.

### Gonadotropin profile in transgenic mice during the estrous cycle

Serum FSH and LH levels were also determined in the animals used in Figure 6. In addition, wild-type female mice of comparable age were sacrificed to cover the different stages of the estrous cycle. Vaginal cytology from the wild-type mice was compared to the transgenic and arranged in an order that would follow a progressive cycle. The wild-type mice displayed the characteristic preovulatory LH surge during proestrus (Figure 7). The transgenic mice also displayed an LH surge during proestrus, although the surge appeared slightly earlier during late diestrus (Figure 7). However, more drastically was an uncharacteristic secondary surge that appeared during late estrus into metestrus.

**Figure 7.**
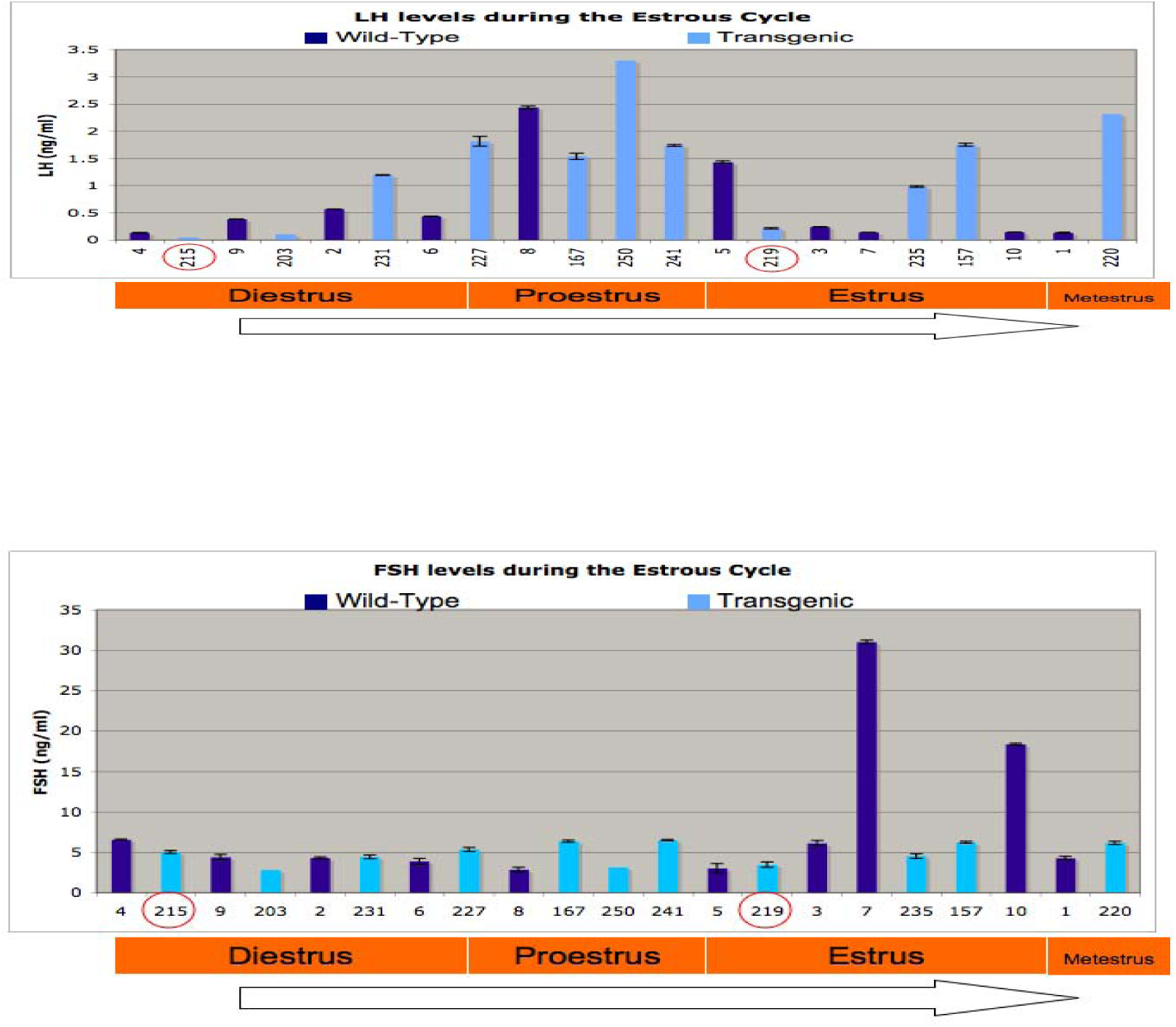
LH and FSH levels during the estrous cycle in transgenic mice compared to wild-type mice. Serum samples were collected from transgenic and wild type mice. Circled are the identification numbers of the transgenic mice that displayed high levels of transgene expression. The stage of the estrous cycle was determined by vaginal cytology. Numbers 1-10 correspond to the wild-type ID number of related age. LH and FSH were measured in duplicates when possible. Error bars are s.e.m.

FSH levels were also measured in these samples, although this piece of data was less conclusive. Although an FSH surge was detected in wild-type animals, this surge occurred during estrus when it is normally detected during proestrus (Figure 7). Surprisingly, there was no surge in serum FSH level in the transgenic mice throughout the examined stages of the estrous cycle (Figure 7).

### Microarray analysis on the ovarian specific ICER transgenic mice

We have shown that selective expression of ICER in mouse ovaries results in an enhanced ovulation rate. Our findings are particularly surprising since Crem null mice do not display obvious female reproductive abnormalities [17,18]. By specifically altering the expression of one CREM isoform, ICER, we may have altered a potential equilibrium existing between the transcriptional activators and repressors in the ovaries, which ultimately lead to this unexpected phenotype. Whereas in the Crem null mice, the equilibrium is cancelled without an obvious effect. This data strongly suggests that ICER plays a fundamental role in the ovulatory process and unraveling the underlying mechanisms of action will help elucidate the role of ICER during ovulation, which are currently being investigated.

Our transgenic mice provide an ideal model for performing in vivo microarray analysis. Presumably, investigating gene regulation differences between transgenic and wild-type, when the transgene is induced, may help determine the overall gene variations involved and the mechanisms by which these animals become hyper-sensitive to ovulatory cues. Therefore, a global gene study between transgenic and wild-type mice was conducted using the Affymetrix GeneChip technology, employing the Mouse Genome 430A 2.0 Array, which contains approximately 14,000 well-characterized mouse genes.

The total RNA extracted from the ovaries of immature transgenic and wild-type mice subjected to exogenous gonadotropin regimen (48hr PMSG + 24hr hCG) used in Figure 2C were utilized for DNA microarray analysis. These samples displayed high levels of the FLAG-ICER protein and RNA in the transgenic ovaries (Figure 2C and 2D), whereas endogenous ICER was not detected at this time point (Figure 3A). All the subsequent technical procedures, including quality control, concentration measurement of RNA, cDNA synthesis and biotin labeling of cRNA, hybridization, and scanning of the arrays, were done at CINJ Core Expression Array Facility of Robert Wood Johnson Medical School (New Brunswick, NJ). Each array was hybridized with cRNA derived from the total RNA sample of either transgenic or wild-type mice (total 2 chips were used in this study).

Expression data from the experimental array (48hr PMSG/ 24hr hCG treated mice) was compared with the non-transgenic control (48hr PMSG/ 24hr hCG wild-type littermate) data. Genes differentially expressed in ovaries of FLAG-ICER mice as a result of superovulation were identified and the changes in gene expression were quantified. All of the candidate transcripts reported in this investigation, passed several selection criteria for stringency purposes. Genes that were significantly regulated 2-fold or more across all the pair-wise comparisons, were considered. Transcripts found to be down-regulated in transgenic mice after superovulation regimen are shown in Table 2. Most of the genes within this list are involved in cytoskeletal organization, muscle contraction and cell adhesion.

**TABLE 2.** Number of transcripts repressed or induced in the ovaries of FLAG-ICER transgenic mice after 48 hr PMSG + 24 hr hCG treatment.

Repressed genes
| Probe Set ID | Gene Title | Gene Symbol | Chromosomal Location |
| --- | --- | --- | --- |
| 1415927_at | actin, alpha, cardiac | Actc1 | 2 E4\|2 64.0 cM |
| 1417023_a_at | fatty acid binding protein 4, adipocyte | Fabp4 | 3 A1\|3 13.9 cM |
| 1417872_at | four and a half LIM domains 1 | Fhl1 | X A6-A7.1 |
| 1417917_at | calponin 1 | Cnn1 | 9 A2-A4 |
| 1418457_at | chemokine (C-X-C motif) ligand 14 | Cxcl14 | 13 B1 |
| 1419100_at | serine (or cysteine) peptidase inhibitor, clade A, member 3N | Serpina3n | 12 F1 |
| 1422340_a_at | actin, gamma 2, smooth muscle, enteric | Actg2 | 6 C3\|6 34.95 cM |
| 1422651_at | adiponectin, C1Q and collagen domain containing | Adipoq |  |
| 1423259_at | inhibitor of DNA binding 4 | Id4 | 13 B\|13 31.0 cM |
| 1423606_at | periostin, osteoblast specific factor | Postn | 3 C |
| 1423669_at | procollagen, type I, alpha 1 | Col1a1 | 11 D\|11 56.0 cM |
| 1423954_at | complement component 3 | C3 | 17 E1-E3\|17 34.3 cM |
| 1424967_x_at | troponin T2, cardiac | Tnnt2 | 1 E4\|1 60.0 cM |
| 1425028_a_at | tropomyosin 2, beta | Tpm2 | 4 B1 |
| 1425506_at | myosin, light polypeptide kinase | Mylk | 16 B3 |
| 1425519_a_at | Ia-associated invariant chain | Ii | 18 E1\|18 32.0 cM |
| 1425546_a_at | transferrin | Trf | 9 F1-F3\|9 56.0 cM |
| 1425811_a_at | cysteine and glycine-rich protein 1 | Csrp1 | 1 E4 |
| 1426731_at | desmin | Des | 1 C3\|1 41.1 cM |
| 1448734_at | ceruloplasmin | Cp | 3 D |
| 1448962_at | myosin, heavy polypeptide 11, smooth muscle | Myh11 | 16 A1\|16 5.0 cM |
| 1449178_at | PDZ and LIM domain 3 | Pdlim3 | 8 B1.1 |
| 1449434_at | carbonic anhydrase 3 | Car3 | 3 A2\|3 11.7 cM |
| 1451342_at | spondin 1, (f-spondin) extracellular matrix protein | Spon1 | 7 F1 |
| 1452670_at | myosin, light polypeptide 9, regulatory | Myl9 | 2 H1 |
| 1460187_at | secreted frizzled-related sequence protein 1 | Sfrp1 | 8 A2\|8 9.5 cM |

Induced genes
| Probe Set ID | Gene Title | Gene Symbol | Chromosomal Location |
| --- | --- | --- | --- |
| 1417017_at | cytochrome P450, family 17, subfamily a, polypeptide 1 | Cyp17a1 | 19 C3\|19 46.0 cM |
| 1422864_at | runt related transcription factor 1 | Runx1 | 16 C4\|16 62.2 cM |
| 1449920_at | cytochrome P450, family 19, subfamily a, polypeptide 1 | Cyp19a1 | 9 A5.3\|9 31.0 cM |
| 1450500_at | U2AF homology motif (UHM) kinase 1 | Uhmk1 | 1 H2 |

Wild-type and transgenic RNA was used for microarray analysis using Affymetrix Mouse Genome 430A 2.0 array representing approximately 14,000 well-characterized mouse genes. These transcripts are down-regulated or up-regulated, 2-fold or more, using pairwise comparison analyses. Experimental data represent n=1.

Transcripts found to be up-regulated in the ovaries of transgenic mice compared to wild type are shown in Table 2. Importantly, CYP19A1 has shown to play a role in fertility as the rate-limiting enzyme in the conversion of androgen to estrogen and has shown to be negatively regulated by ICER [8]. Also, worth noting, CYP17A1, a steroid 17-alpha-hydroxylase and a steroid 17, 20-lyase participating in both glucocorticoid and estrogen biosynthesis. To confirm the responsiveness of these genes to cAMP, we utilized the CREB Target Gene Database to validate the microarray data and to determine whether the genes of the corresponding transcripts contained a CRE in their promoter region [19]. We found that the majority of the regulated transcripts were shown to contain a predicted CRE within their gene and have been reported cAMP-regulated according to the CREB Target Gene Database.

### Real-Time PCR validation of the microarray results

We next validated the microarray data by confirming the differential expression pattern by real-time PCR (Figure 8). We selected two genes which were up-regulated (Cyp17a1 and Cyp19a1) and one gene that was down-regulated (Des) to monitor the expression levels. We also measured the levels of cyclin D2 and INHA transcripts. Relative quantification real-time PCR was performed to determine changes in target gene expression within the transgenic sample relative to the wild-type littermate used in the DNA microarray analysis. This allows for comparison between the initial levels of template in each sample, without requiring the exact copy number of the templates. As shown in Figure 8, the expression pattern of Cyp17a1, Cyp19a1 and Des determined by microarray, were confirmed by real-time PCR. Moreover, the levels of cyclin D2 transcript remained unchanged. Surprisingly, the expression of INHA was found to be around 5-fold higher in the transgenic mice.

**Figure 8.**
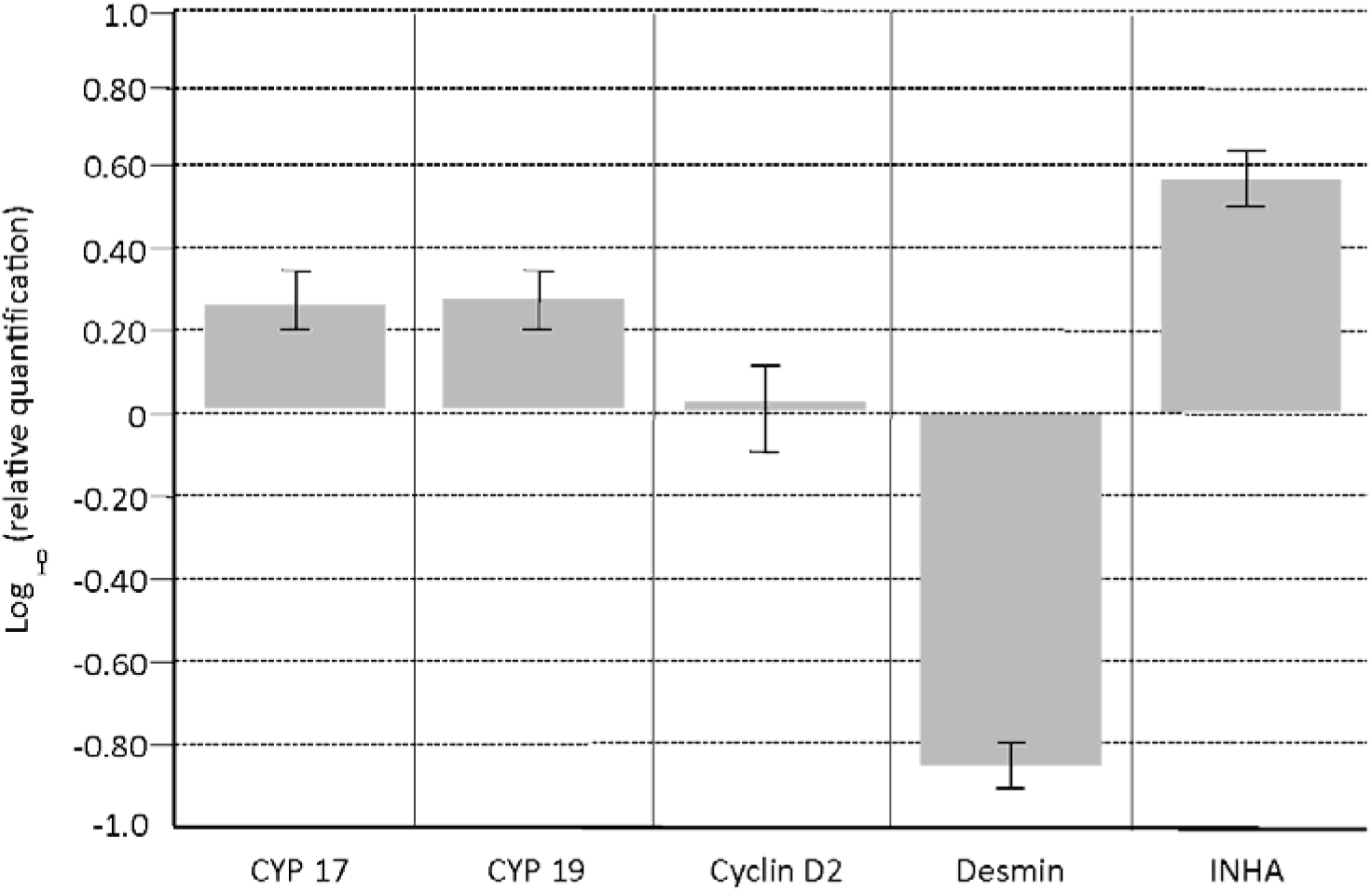
Comparative gene expression using quantitative real-time PCR of several genes in the ovaries of FLAG-ICER using wild-type ovaries as the calibrator. A relative quantification assay was used to determine the changes in expression of Cyp17a1, Cyp19a1, Cyclin D2, desmin and inhibin alpha in transgenic samples used for DNA microarray analysis relative to the Wild-type sample. Microarray analysis showed Cyp17a and Cyp19a to be up-regulated in the transgenic samples (Table 3) whereas Desmin was shown to be down-regulated (Table 2). Cyclin D2 was measured as a negative control, since the expression levels was unchanged between transgenic and wild-type. Samples were reverse-transcribed and the cDNA was subjected to real-time PCR quantification. All gene expression levels were normalized to GAPDH expression levels and expressed as a fold change from the wild-type. Fold-expression changes were calculated using the delta-delta Ct method. Each sample was the mean of triplicate real-time PCR experiments. The error bars displays the calculated maximum (RQMax) and minimum (RQMin) expression levels that represent standard errors of the mean expression level (RQ value).

## DISCUSSION

Currently an ovarian-specific ICER-null mouse model is not available to fully validate the role of ICER during folliculogenesis. *Crem*-mutant mice have been generated resulting in male sterility due to postmeiotic arrest at the first step of spermatogenesis, whereas the female mice were reported to be fertile [17,18]; although detailed studies of the reproductive processes which occur in the females have not yet been reported. Judging by the apparent pleiotropic function of CREM, such a specific phenotype is surprising. Since the homozygous mutant mice lacked activators and repressors such as CREM and ICER, we suspect that the balance mechanism controlling CREM-mediated gene expression in *Crem*-null mice might be affected in two opposite directions. Which would result in a mutual cancellation without apparent phenotypic differences in female mice.

ICER and CREM proteins are both detected in the ovaries hence, the generation of an ovarian-specific either ICER-null mice or ICER transgenic animals will aid in answering these questions by disrupting this presumed balance. In accordance with this postulate, we successfully generated ovarian-specific ICER transgenic mice in order to disrupt this potential balance. Exogenous levels of FLAG-ICER transcript and protein were highly detected in the ovaries of the transgenic mice when subjected to gonadotropin treatment (Figure 2). Immunohistochemistry analysis of the ovaries of the transgenic mice treated with PMSG and hCG showed that the spatial expression of FLAG-ICER is similar to the endogenous expression mainly localized to the granulosa cells of antral follicles as well as in the luteal cells of the developing corpus luteum (Figure 3B). However, the temporal pattern of expression of exogenous ICER in the transgenic mice strongly deviated from the previously reported expression pattern of endogenous ICER [11]. Ovaries from immature rats subjected to exogenous gonadotropin treatment normally express ICER within 4hr of hCG treatment and declines after 7hr; whereas immature transgenic female mice display induced levels of exogenous ICER 48hr after of PMSG treatment with enhanced and sustained expression with the subsequent treatment of hCG for 24hrs. Moreover in mature, estrous cycling, transgenic mice the temporal pattern of expression of the transgene displayed a biphasic expression during diestrus and proestrus (Figure 6), contrast to observed expression pattern of endogenous ICER, which is normally expressed only during proestrus [11]. Surprisingly, these transgenic mice display biphasic surge in LH, during proesterus and in late estrus/early mettestrus (Figure 6). The consequence in the altered temporal expression of ICER may account for the altered levels of LH detected in the transgenic mice.

We have also shown that the selective expression of ICER in the ovaries of transgenic mice results in enhanced ovulation rate. Our finding is particularly surprising since CREM/ICER null mice, as mentioned above, did not display obvious female reproductive abnormalities [17,18]. The physiological consequence manifested in the ICER transgenic mice raises the insisting question: Why does the ablation of ICER in the CREM null mice produce no change in phenotype? Perhaps by specifically altering the expression of one of the CREM isoforms, ICER, we negated any potential equilibrium that may have existed between the transcriptional activators and repressors in the ovaries.

Nevertheless, this data strongly suggests that ICER potentially regulates ovulation. The underlying mechanism responsible for this regulation is currently being investigated. We can hypothesize that ICER may inhibit the degeneration and reabsorption of the ovarian follicle before it reaches maturity and ovulates, a process referred to as follicular atresia. Inhibiting follicular atresia would rescue atretic follicles from their destined fate, resulting in an increase in the number of follicles that mature and ovulate. Another possibility is that the follicles become more receptive to the effects of gonadotropins, potentially through an increase in the quantity of FSH/LH receptors, which would lead to more efficient ovulation events.

ICER has previously been implicated in the transcriptional repression of three FSH responsive genes, *Inha*, *Cyp19a1 and cyclin D2.* The subsequent alteration in the temporal expression of ICER in the transgenic mice prior to an ovulatory cue could essentially mimic the phenotype manifested in mice with null mutations in *Inha* or *Cyp19a1*. However, *Inha* null mice display high levels of serum FSH and the peptide hormone activin, which enhances FSH synthesis and secretion, ultimately resulting in infertility in female mice before succumbing to granulosa cell tumors [20, 21]. Furthermore, female mice with null mutations on *Cyp19a1* manifest an increase in circulating testosterone levels with subsequent alterations in FSH and LH levels, thereby resulting in infertility due to disruption in folliculogenesis [22]. Whereas null mutations in cyclin D2 has been shown to lead to infertility due to lack of granulosa cell proliferation resulting in the disruption of folliculogenesis [14]. Thus, we assumed that our transgenic animals would be infertile based on our previous data showing ICER regulation of cyclin D2 [9]. However, the ICER transgenic mice model displayed enhanced ovulatory rates in response to stimulatory effects of the gonadotropins. Several other transgenic mouse models with specific ovarian follicle phenotypes has been described [23]. To date, this is the first transgenic mouse model of its kind. As surprising as our findings might be, it is important to state that our adult transgenic mice do not express the transgene throughout the entire length of the estrous cycle, nor do they display a gross over-expression of ICER levels.

The 2.5kb portion of the *Inha* gene contains a CRE which has been proposed as the site of transcriptional activation by CREB in response to FSH, followed by the transcription repression during the LH surge as a result of ICER induction [7, 11]. In the transgenic animals, FLAG-ICER inducibility in mature mice may be followed by an auto-regulation of the transgene. This would account for the lack of sustained transgene expression; until the preovulatory LH surge results in the expression of the transgene again. Although theoretically the *Inha* gene is normally repressed as a result of the LH surge, the 2.4kb region we are utilizing may not contain the regulatory elements required for LH repression. This effect is more drastic in mice where hCG treatment in PMSG primed mice results in a prolonged expression of the transgene. It is possible that FLAG-ICER is mediating gene regulation at certain stages of the estrous cycle, with alternating relief of the regulation upon auto-regulating its own expression. This intricate expression system may perhaps promote and enhance response to the ovulatory cues, thereby resulting in hyperovulation.

Investigating the differences in gene regulation between transgenic and wild-type mice prior to ovulation may identify a mechanism by which these animals become hyper-sensitive to ovulatory cues. These animals provide a suitable model to perform *in vivo* microarray analysis, allowing us to unravel the underlying gene regulation responsible for the observed hyperovulation. Consequently, these transgenic mice constitute a unique model to study fertility and the physiological cues that would enable us to control the rate of ovulation.

## ACKNOWLEDGMENTS

We thank M. Meyenhofer for technical assistance in preparing the tissue sections, L. Ampow for assistance in the preparation of this manuscript and for technical assistance with the immunostaining and Cory Haluska for critical reading of the manuscript.

